# ProFam: Open-Source Protein Family Language Modelling for Fitness Prediction and Design

**DOI:** 10.64898/2025.12.19.695431

**Authors:** Jude Wells, Alex Hawkins Hooker, Micha Livne, Weining Lin, David Miller, Christian Dallago, Nicola Bordin, Brooks Paige, Burkhard Rost, Christine Orengo, Michael Heinzinger

**Author notes:** Equal contribution (Jude Wells, Alex Hawkins Hooker).

## Abstract

Protein language models have become essential tools for engineering novel functional proteins. The emerging paradigm of family-based language models makes use of homologous sequences to steer protein design and enhance zero-shot fitness prediction, by imbuing models with an ability to explicitly reason over evolutionary context. To provide an open foundation for this modelling approach, we introduce *ProFam-1*, a 251M-parameter autoregressive protein family language model (pfLM) trained with next-token prediction on millions of protein families represented as concatenated, unaligned sets of sequences. *ProFam-1* is competitive with state-of-the-art models on the ProteinGym zero-shot fitness prediction benchmark, achieving Spearman correlations of 0.47 for substitutions and 0.53 for indels. For homology-guided generation, *ProFam-1* generates diverse sequences with predicted structural similarity, while preserving residue conservation and covariance patterns. All of ProFam’s training and inference pipelines, together with our curated, large-scale training dataset *ProFam Atlas*, are released fully open source, lowering the barrier to future method development.

## 1 Introduction

Protein language models (pLMs) distill statistical patterns from vast sequence databases to predict protein properties [1, 2], score variants [3, 4], and guide engineering [5, 6]. Dominant pre-training strategies include masked language models (MLMs) [1, 2], sequence diffusion models [6–8], and autoregressive pLMs [9–11]. While MLM likelihoods excel at ranking point mutations, they are ill-suited for scoring the relative fitness of insertions, deletions, and distant sequence variants. Conversely, autoregressive models offer a principled framework by decomposing the joint likelihood into a series of conditional probabilities, providing a likelihood score that is efficiently computed and that naturally accommodates insertions and deletions. Moreover, sampling from a learned autoregressively decomposed probability distribution supports *de novo* generation without any fixed constraint on sequence length or alignment to a given seed sequence. However, unconditional autoregressive generation has limited practical utility, as it samples from the entire protein sequence space rather than targeting specific protein families or functions [12].

Early strategies for steering generative models relied on explicit functional labels, such as enzyme commission numbers [13], or fine-tuning on specific protein families [14]. Recent approaches instead condition directly on evolutionary context provided by sets of homologous sequences. This paradigm includes MSA-based MLMs [15, 16] and generative models [7, 17, 18] as well as alignment-free autoregressive models trained on concatenated sequences [19–21]. Unlike functional tags, conditioning on homologs guides generation via evolutionary constraints without requiring rigid functional priors. For scoring, the family context in the prompt provides evolutionary information that improves fitness prediction [16, 19, 21]. For sequence generation, family conditioning directs sampling toward a design goal via in-context exemplars, without the need for any additional model fine-tuning.

Here, we introduce ProFam, a suite of data, weights, and code for the training and inference of autoregressive protein family language models (pfLMs). First, we introduce a curated, large-scale and openly accessible training corpus called *ProFam Atlas*, incorporating single and multi-domain protein sets derived from sequence-, structure- and function-level relationships. To exploit this dataset, we introduce a 251M parameter autoregressive Transformer model, *ProFam-1*, trained on ProFam Atlas. We provide open-source training and inference code together with model weights.

In terms of capabilities, ProFam-1 achieves performance competitive with state-of-the-art sequence-only methods on the ProteinGym zero-shot fitness prediction benchmark. Motivated by use-cases in evolution-based functional protein design, we also demonstrate ProFam’s sequence generation capacities via a series of *in silico* evaluations measuring the ability of the model to produce sequences recapitulating features of the distribution of natural protein families. To maximise the signal available to the model via the evolutionary context in both settings, we experiment with sequence prompt ensembling schemes, finding that they improve both the diversity of the model’s generations and its ability to follow evolutionary constraints. Finally, we explore ProFam’s potential use in augmenting available evolutionary signal by generating synthetic MSAs for CASP targets [22], finding that they improve ColabFold-based structure prediction over single-sequence inputs [23].

## 2 The ProFam Atlas Dataset

We introduce a curated, large-scale dataset of protein sequences grouped by varying definitions of *family* (Figure 1A). These family definitions include structurally-related multi-domain proteins in the AlphaFold Database (AFDB) [24], functionally consistent structure-derived domain families in The Encyclopedia of Domains (TED) [25], and deep Multiple Sequence Alignments (MSAs) from the maximally diverse 270k-representative subset of the OpenFold OpenProteinSet [26]. Each of these data sources contains a set of families, where each family consists of a set of protein sequences. In addition to families, the ProFam Atlas dataset also contains single sequences from UniRef90 to increase the coverage of protein sequence space. The details of individual data sources are provided in Appendix A. We hypothesise that allowing for various definitions of family in training data encourages models to learn from varying types of homology signal at training time, thereby increasing their flexibility at inference time by enabling the support of various types of evolutionary context as input prompts. In total, the ProFam Atlas contains almost 40 million protein families and around 481M protein sequences (not accounting for redundancy between families). This resource provides a uniquely comprehensive open foundation for training and evaluating family-based pLMs.

**Figure 1.**
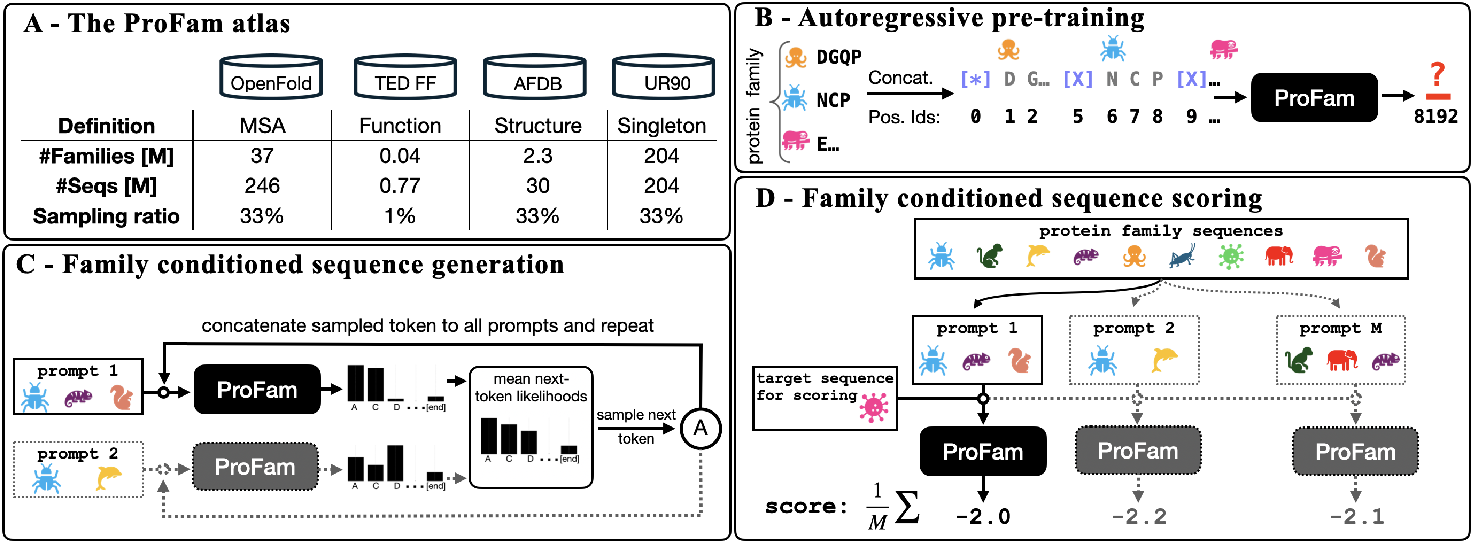
The ProFam Atlas & ProFam-1 Training and Inference. The ProFam Atlas (Panel A) consists of protein families derived from the AlphaFold-Database (AFDB), TED FunFams (FF), and the OpenFold OpenProteinSet as well as individual proteins from UniRef90. ProFam-1 (Panel B) is autoregressively pre-trained on protein families by sampling at different ratios from the data sources contained in ProFam Atlas. For each sample, unaligned protein sequences within one family are concatenated using special separator tokens (depicted here as [X]) up to a maximum of 8192 tokens/amino acids per family. During inference, ProFam-1 can leverage evolutionary information from homologs to either guide generation towards proteins with related function and structure (Panel C) or score variants (Panel D). Instead of expanding the model’s input to ultra-long context, both scoring and generation can be improved by ensembling large families into multiple prompts (indicated by dashed lines and gray coloring in panels C and D), each processed separately, and we average either the next-token probabilities (generation) or the mean of the per-token log-likelihoods from the variant of interest (variant scoring).

## 3 The ProFam-1 Model

ProFam-1 is an autoregressive Transformer language model built on the Llama architecture [27] with 16 layers, 1024 hidden dimensions, and approximately 251 million parameters. ProFam-1 is trained on a mix of single sequences and unaligned, concatenated protein sequences (up to 8192 tokens) from the ProFam Atlas families. Since many protein families substantially exceed this context window, each family-level training example is constructed by randomly subsampling sequences from the family, until the 8192-token budget is reached. Batches consist of sets of such training examples, sampled uniformly (33% each) from AFDB, OpenProteinSet and UniRef90 but with lower probability from TED FunFams to avoid oversampling of the relatively few families in this set (Appendix Table A.4). We use one token per amino acid as well as special tokens to mark the start of a document and the end of an individual protein sequence. For positional encoding, we use the default Llama RoPE encoding scheme [28] for positions within the concatenated sequence-of-sequences string. This design preserves both within-sequence and between-sequence positional information, while avoiding the complexity of non-standard Transformer architectures used in prior works [19]. During training, we randomized the order of sequences within each document to encourage invariance with respect to sequence order. With this setup, the model was trained for 592,619 steps covering 236 billion tokens, for a total of approximately 14 days on four NVIDIA A100 GPUs.

As an autoregressive protein family language model, ProFam-1 represents a probability distribution over the sequences in a family [19]. This probability distribution can be used for both *sequence scoring*, i.e. evaluating the likelihood of a protein sequence given a (possibly empty) set of sequences belonging to a given family, and *sequence generation*, i.e. sampling novel sequences belonging to the same family as the sequences in the conditioning set. In practice, in both cases, a ‘prompt’ of concatenated sequences of up to 8192 tokens in total length is constructed and passed to the model. There is considerable freedom in how this prompt is constructed for a given family, with the choice of prompt determining the conditional distribution over the family represented by the model. We explore some of these details in this paper, and describe relevant methods in the following sections.

## 4 Inference-Time Strategies for Scoring and Sampling Sequences

### 4.1 Sequence Scoring

Autoregressive protein *family* models, such as ProFam-1 and PoET [19], predict each amino acid in a variant sequence while conditioning on (i) the already-generated prefix of the variant and (ii) a *prompt* consisting of homologous family sequences. Let *S* = (*s*_1_, …, *s*_*L*_) denote a single protein variant of length *L*, to which we append a special end-of-sequence token *s*_*L*+1_. Let ℛ denote the set of all available family members (homologous sequences) for the corresponding protein family. We write 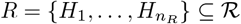 for a (possibly subsampled) prompt set of homologs, where each *H*_*j*_ is a protein sequence.

We score a variant using its *average per-token log-probability* under the model conditioned on *R*, including the prediction of the end-of-sequence token:

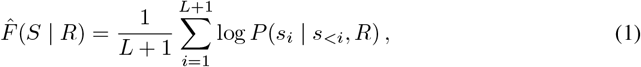

where *s*_*<i*_ := (*s*_1_, …, *s*_*i*−1_) denotes the variant amino acids up to (but not including) position *i*. Conditioning on evolutionarily related sequences allows family-based models to leverage evolutionary information when estimating variant effects, which has been shown to improve fitness prediction accuracy compared to single-sequence models in both unsupervised [19] and supervised [29] settings. To encourage sequence diversity among the sequences sampled for the prompt, we use the same sequence weighting scheme used by PoET [19]. This approach is described in detail in Appendix D.

### 4.2 Evolutionary Context Ensembling

Analogous to the concept of test-time-scaling in LLMs [30], which achieves better downstream performance via additional inference-time forward passes, family-based protein language models can exploit ensembling over multiple distinct subsampled input sequence sets to improve performance [15, 19, 29]. Previous works have demonstrated the effectiveness of this strategy in improving zero-shot and supervised fitness prediction. We build on these methods by explicitly exploring the scaling behaviour of input context ensembles in the fitness scoring setting, as well as extending the strategy to generation via a strategy we refer to as ensemble sampling.

#### Ensemble Scoring

For scoring, we sample *M* prompts independently from ℛ, denoted 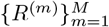 with *R* ^(*m*)^ ⊆ ℛ. We then average the per-prompt log-probability scores:

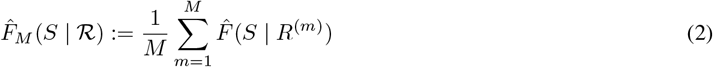

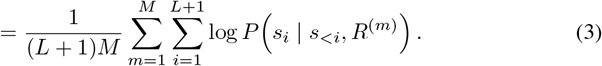

#### Ensemble Sampling

Our ensemble sampling strategy aggregates prompts by averaging their next-token probability distributions (softmax outputs) at each step and sampling the next token from the resulting ensemble distribution. In contrast, scoring averages log-probabilities of the observed tokens across prompts.

Formally, at generation step *i*, for each prompt *R* ^(*m*)^ (*m* ∈ { 1, …, *M*}), let 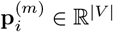 denote the model’s predicted probability distribution over the vocabulary *V*, computed given the prefix *s*_*<i*_ and prompt *R* ^(*m*)^. The ensemble distribution is given by the mean:

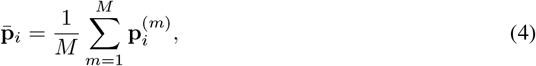

from which we sample 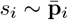, then append the sampled token to the prefix. For simplicity, we omit top-*p* (nucleus sampling) and take temperature 1 by default. Additional sampling hyperparameters are provided in Appendix E.1. When evaluating ensemble sequence generation with ProFam-1, we used *M* = 8 independently sampled prompts.

## 5 Results

In the following sections, we first describe ProFam-1’s zero-shot fitness prediction capabilities as evaluated on the widely-used ProteinGym benchmark. We then assess sequence generation across a suite of *in silico* tasks relevant for functional protein design. Details of all benchmark settings are provided in Appendix C.

### 5.1 Improved Zero-Shot Fitness Prediction Using Test-Time-Scaling

We assess the correlation of evolutionary-context-aware ProFam-1 likelihoods with protein fitness using deep-mutational scanning (DMS) data from ProteinGym [31]. Prompt ensembling (Section 4.2) transforms ProFam from an average predictor (Spearman 0.42) into one that is competitive with state-of-the-art (Spearman 0.47) (Fig. 2). We observe that aggregating log-likelihoods across diverse prompts steadily raises Spearman correlation, with larger ensembles closing much of the gap to the very best zero-shot methods (Fig. 2, panel C). Second, fitness performance is not monotonically increasing with likelihood. Across assays, pushing the average variant log-likelihood too high often reduces Spearman; performance peaks in a mid-log-likelihood regime (roughly −1.8 to −1.1, often near −1.3). When using few prompts/ensembles, tuning the individual prompt(s) to target this likelihood range improves correlation; with larger ensembles, the benefit diminishes as ensemble diversity dominates (fig. 2C).

**Figure 2.**
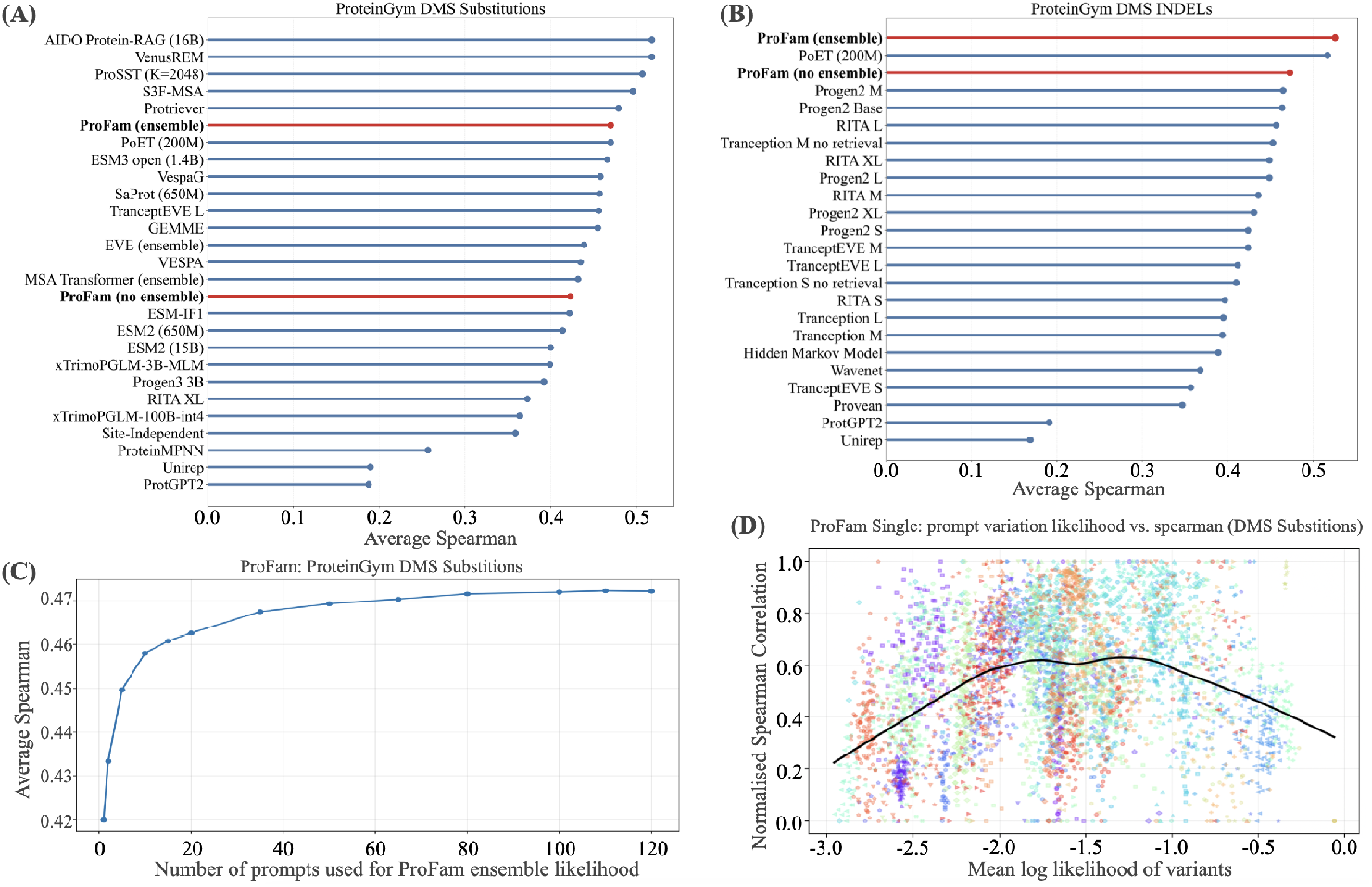
Test-Time-Scaling Improves ProteinGym Zero-shot Performance. In Panels A and B we show the average Spearman correlation between model predictions and experimental fitness scores across 217 and 65 deep mutational scanning experiments. Panel A shows substitutions (all assessed variants must be the same sequence length as the wild-type), while panel B shows performance for insertions and deletions. Panel C shows how ProFam’s performance increases with additional prompts used for ensembling the likelihood score. In Panel D we plot results for 50 randomly selected ProteinGym DMS assays (no indels). Each assay is represented with a color-marker combination. For each assay, we show up to 150 results from different prompts (different context sequences) with the average log-likelihood of the variants on the x-axis and the normalised Spearman correlation between the variant log-likelihood and the experimentally measured fitness score on the y-axis. We normalise so that the Spearman score has a minimum value of 0 and a maximum value of 1 across all prompts. We see that the ‘best’ prompts are those which yield a variant log-likelihood in the range −1.8 to −1.1.

### 5.2 Prompt Engineering for Improved Fitness Prediction

As shown previously [32, 33], higher average log-likelihood does not monotonically translate into better zero-shot fitness prediction. As shown in Figure 2 D, performance improves as the variant log-likelihood increases up to roughly −1.8, but beyond approximately −1.1 the trend reverses: prompts that further raise likelihood tend to yield lower Spearman correlations with experimental fitness. Figure 2 D shows correlations scaled so that values for each assay span the [0, 1] range. This highlights the trend, but not the scale. In contrast, Appendix Figure 7 displays the unnormalised Spearman values for each prompt-assay combination for 25 assays, with one point per prompt (i.e., per subsample of the family). Two patterns emerge. First, the attainable Spearman correlation varies widely across prompts for the same assay: the gap between the best and worst prompt commonly exceeds 0.3. Second, the average variant log-likelihood also spans a broad range, and the optimal likelihood differs by assay.

### 5.3 ProFam-1 Generations Recapitulate the Statistics of Natural Families

ProFam-1 demonstrates a consistent advantage over baselines in capturing key family statistics in EC families, particularly when generating from a single sequence. Under single-sequence conditioning, ProFam-1’s synthetic MSAs achieve higher conservation correlation and lower KL divergence (natural → synthetic MSA) in position-specific amino acid distributions (Figure 3a–b) compared to PoET. With multi-sequence conditioning, ProFam-1 generates a notably higher proportion of sequences within an acceptable length range (95.1% vs. PoET’s 83.8%; Fig. 3d) and exhibits lower KL divergence when mean sequence identity is less than 80 (Fig. 3e). As expected, conditioning on multiple sequences generally reduces KL divergence compared to single sequence conditioning (Fig. 3e vs. 3b). We also assessed ProFam-1’s ensemble mode (Section 4.2). Ensemble generation yields sequences with lower average pairwise identity (PID) to natural homologs and typically achieves lower KL divergence at comparable PID levels (Fig. 3e). This increased diversity, however, slightly reduces the proportion of sequences within the acceptable length range (Fig. 3d).

**Figure 3.**
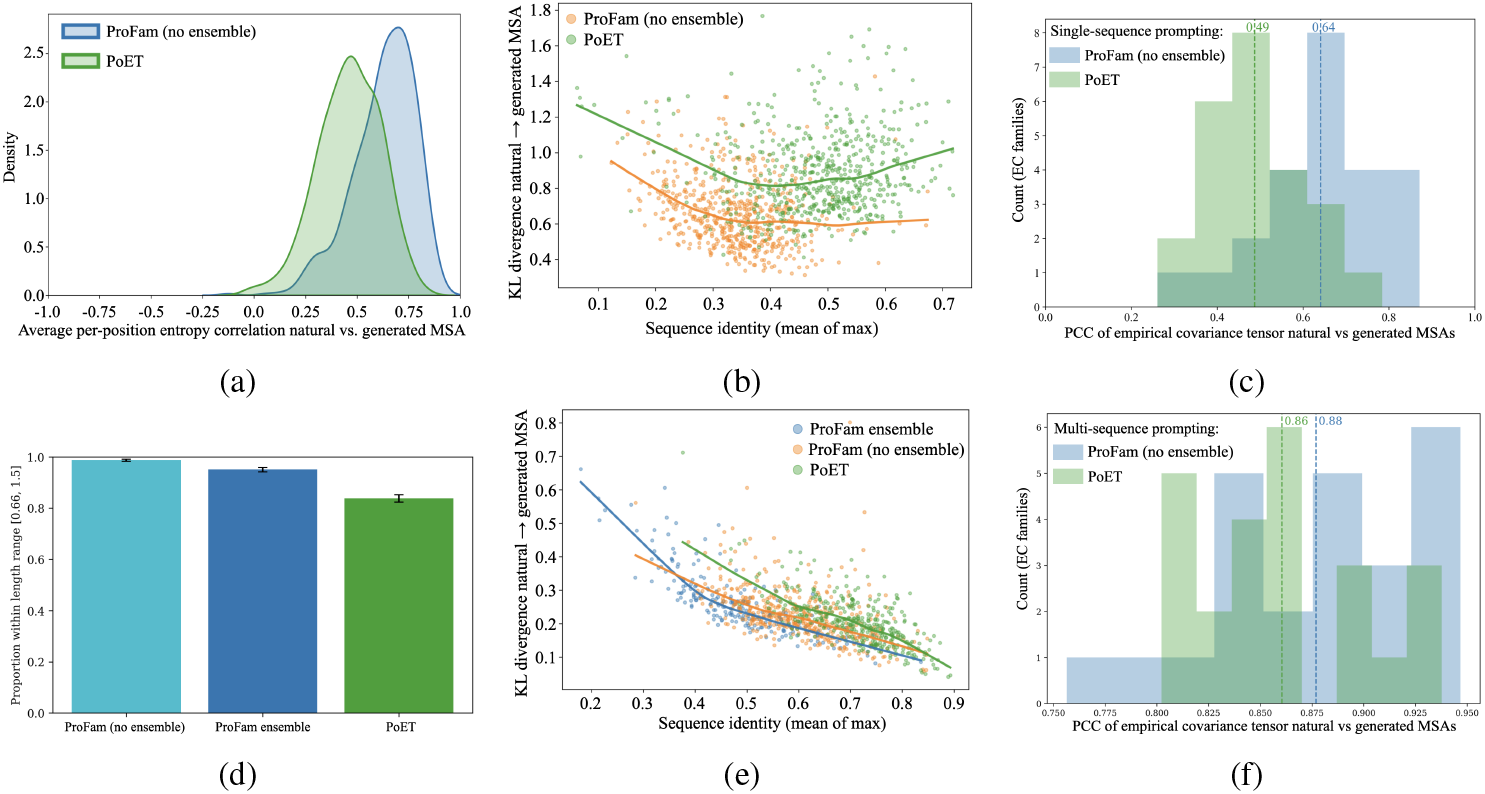
ProFam-1 Captures Family Statistics. We tested how well ProFam-1 and PoET capture perposition conservation correlation (3a) and KL divergence of position-specific amino-acid distributions (3b) under single-sequence EC prompting. Using the same set of families, we show how multi-sequence prompting changes KL divergence (3e) and how many generated sequences have acceptable length (3d). On a reduced set of 24 large EC families, we show that ProFam-1 (and PoET) capture higher-order statistics by correlating covariances extracted from synthetic and natural alignments. The gap between ProFam-1 and PoET is bigger for single-(3c) than for multi-sequence prompting (3f).

Moving beyond single-site statistics, we evaluated the extent to which ProFam-1 captures higher-order dependencies as quantified by the Pearson Correlation Coefficient (PCC) of covariances derived from natural and synthetic MSAs. For this, we used the subset of 24 deep EC families, described in Appendix section B.1, generated 1200 proteins conditioning on a single prompt per family and computed covariances as described in Appendix section C.3. By focusing only on positions with sufficient support, we avoided introducing noise from positions that are not well supported in the natural MSA. Following this analysis, we see that both PoET and ProFam-1 learn to extract higher-order correlations from single protein sequences (PCC of 0.49 and 0.64, respectively; 3c which improves notably with multi-sequence prompting (PCC of 0.86 and 0.88, respectively; Fig. 3f).

To quantify the utility of generated sequences, we input ProFam-1 synthetic MSAs (generated by conditioning on the target sequence only) for hard (free-modelling) CASP15/16 targets into ColabFold and measured structure prediction accuracy using lDDT and TM-score against experimental structures (Section C.4). Although synthetic MSAs (lDDT=0.7) did not match the performance of natural MSAs (lDDT=0.9), they improved over single-sequence inputs (lDDT=0.5). Gains in TM-score were more modest, showing only a non-significant upward trend compared to single sequence structure prediction, with natural MSAs remaining superior (Fig. 4).

**Figure 4.**
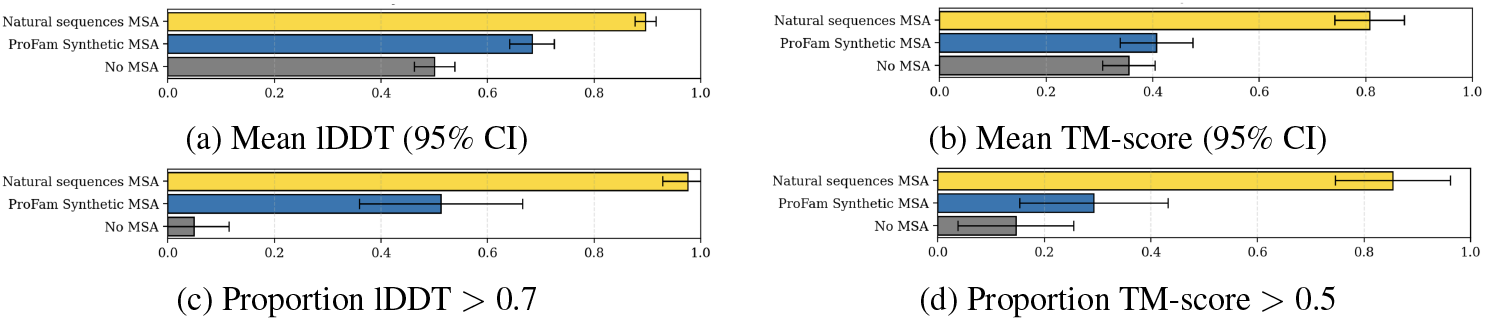
ColabFold performance on targets from CASP15/16 using ProFam-1 synthetic MSAs. We compare single-sequence inputs (no MSA), ProFam-1 synthetic MSAs, and natural MSAs. The plots show the mean (with 95% CI) for lDDT and TM-score, as well as the proportion (with 95% CI) of predictions exceeding standard quality thresholds (lDDT *>* 0.7, TM-score *>* 0.5).

### 5.4 ProFam Generates Diverse Sequences Retaining Family Structure

To evaluate the design capabilities of ProFam-1, we conditioned on homologs within the 128 hold-out families, predicted the structure of the generated sequence and compared it to the AFDB entries of natural proteins within the same family using TM score [34]. Additionally, we report for each generated sequence the mean predicted local difference distance test (plDDT), an AlphaFold confidence metric [35]. When plotting these metrics against the maximum sequence identity across family members, including those sequences that were not sampled in the prompt (Fig. 5a, 5b), we find that ProFam-generated sequences (in both non-ensemble and ensemble mode) have similar TM and pLDDT scores to PoET, with a non-significant trend favouring ProFam when the PID is less than 70 (Fig. 5a). If we restrict our analysis to only consider generated sequences with PID less than 50 to any natural sequence in the family, we observe a modest, non-significant increase in the proportion of ProFam-1 sequences with TM > 0.5 relative to PoET (Fig. 5c).Both ProFam-1 and PoET significantly outperform the random mutation baseline, where random mutations are added to natural sequences at different rates.

**Figure 5.**
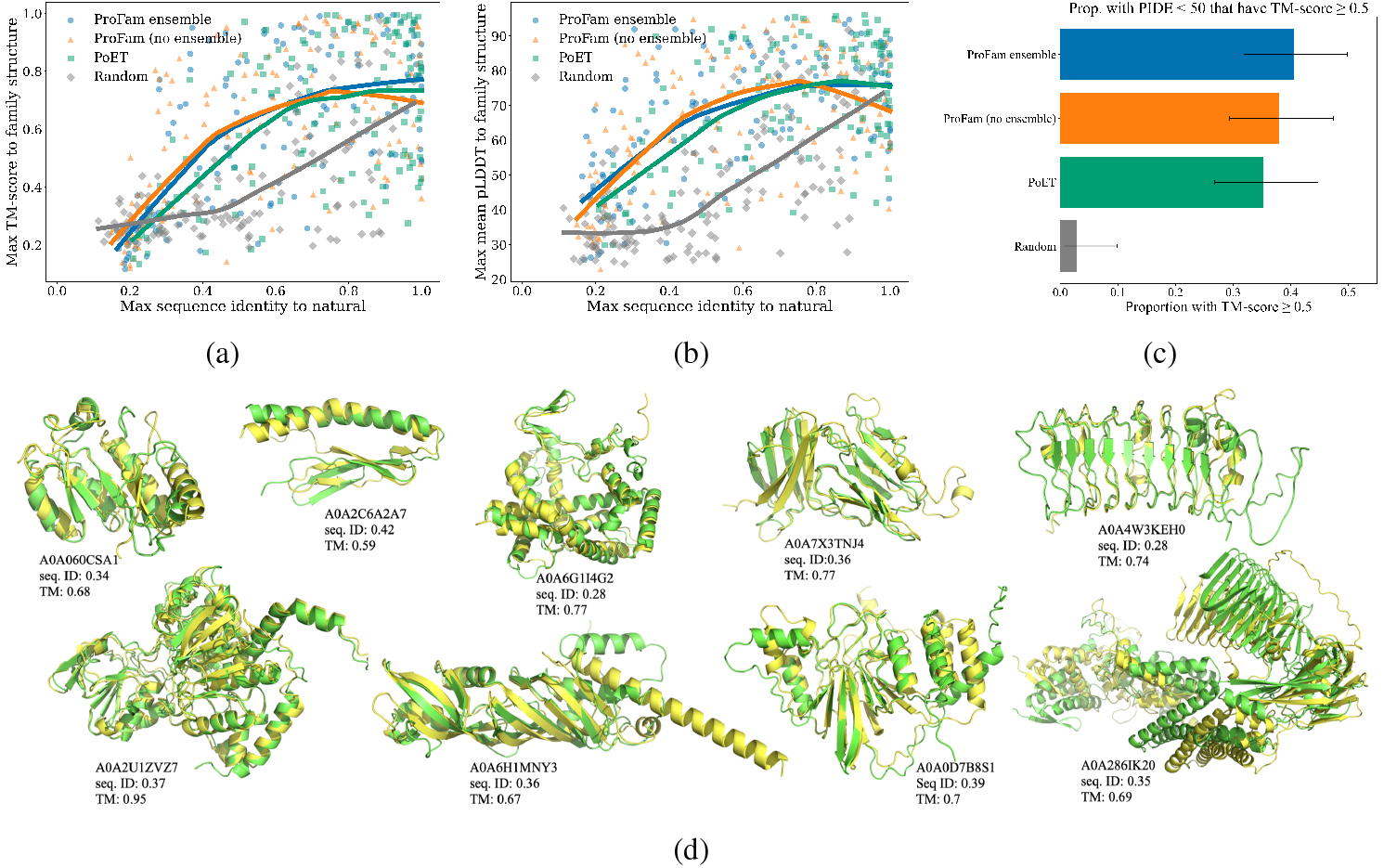
ProFam Generates Diverse Sequences for Given Family Fold. We compare sequences generated by ProFam-1 and PoET when conditioned on sequences from 128 held-out FoldSeek cluster families. We also show results for a random mutation baseline generated by randomly changing amino acids in natural sequences from the family. Structural coherence of the generated sequences was quantified by comparing ColabFold predictions of generated sequences with the family structures (taken from AFDB). 5a shows the relationship between maximum sequence identity and maximum TM-score when comparing the generated sequence with the natural family sequences and structures; similarly, 5b shows this relationship with pLDDT. 5c restricts the analysis to generated sequences with less than 50% maximum sequence identity to any natural family member, we report the proportion with TM-score greater than 0.5 (with 95% CI). 5d shows predicted structures of ProFam-sampled sequences (green) overlaid with the predicted structure of the most similar natural sequence (yellow). For each pair, we report the UniProt ID, sequence identity and TM score. Sampling hyperparameters for PoET and ProFam are listed in Appendix Section E.1

## 6 Conclusion

We present ProFam, a comprehensive open-source framework for protein family language modelling, comprising the 251M parameter ProFam-1 model and the ProFam Atlas: a dataset of nearly 40 million protein families. We show that ProFam-1 achieves competitive performance on zero-shot fitness prediction and demonstrates strong sequence generative capabilities. To validate the latter, we conducted structural consistency checks of generated sequences on 128 FoldSeek clusters from the AFDB and rigorous statistical assessments across 460 EC families. We further demonstrate how information-theoretic comparisons between natural and synthetic MSAs can be used as an in silico benchmarking strategy for assessing how well pfLMs capture family-level sequence statistics, while noting that future work is needed to establish how predictive these metrics are of experimentally measured protein function.

Our results highlight that when conditioned on a single exemplar sequence, ProFam-1 consistently generates sequences with higher fidelity to family conservation statistics compared to PoET. When generating synthetic MSAs from ProFam-1 for structure prediction, we observed a statistically significant improvement in lDDT scores and a non-significant upward trend in TM-scores compared to single-sequence inputs. Crucially, we found that large gains in lDDT and pLDDT often did not correspond to visible improvements in the fold, whereas TM-score improvements were much better correlated with structural quality. While we identified specific cases where the correct fold emerged solely through conditioning on ProFam synthetic MSAs (Appendix Fig. 6), we found no instances where synthetic MSAs yielded better predictions than MSAs constructed from natural homologs. Nonetheless, the ability to imprint realistic coevolution could be exploited to bootstrap MSAs for de novo or sparsely annotated designs where no natural homologs exist.

**Figure 6.**
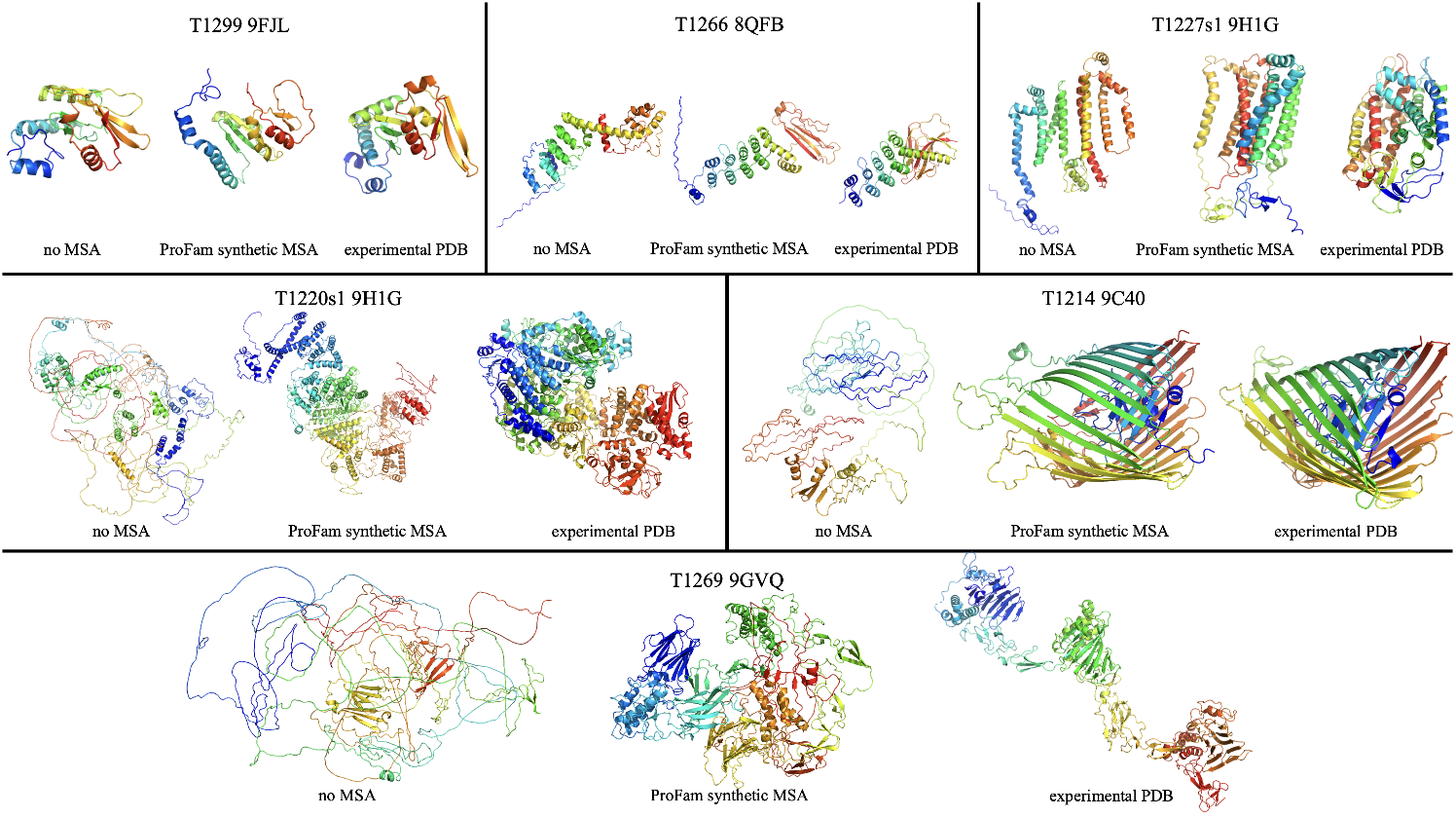
ProFam Synthetic MSAs Improve Structure Prediction for CASP16 targets. AlphaFold2 predicted structure comparison for six CASP16 targets using: no MSA (left), and a ProFam synthetic MSA generated from the target sequence alone (middle). ProFam synthetic MSAs improve over no MSA in 5 of 6 visualised cases (T1266, T1227s1, T1220s1, T1214, T1269) and are worse in one case (T1299); Experimental PDB structures are shown on the right. Predictions using MSAs of real homologous sequences (not shown) remain the best across all targets.

In the domain of zero-shot fitness prediction, we examined the interplay between likelihood and fitness. We observe a “sweet-spot” in average variant log-likelihood (approximately −1.3) where zero-shot performance peaks. Importantly, when randomly sub-sampling a MSA to create single prompts, the choice of evolutionary context can drastically change fitness scoring (Appendix Fig. 7). While trying to leverage this observation by carefully engineering prompts to hit the target likelihood improved performance, we find that large-scale prompt ensembling renders such engineering largely redundant. By aggregating scores over 120 randomly sampled prompts, ProFam-1 achieves fitness prediction performance on ProteinGym competitive with state-of-the-art models (0.47 on substitutions and 0.53 on indels).

**Figure 7.**
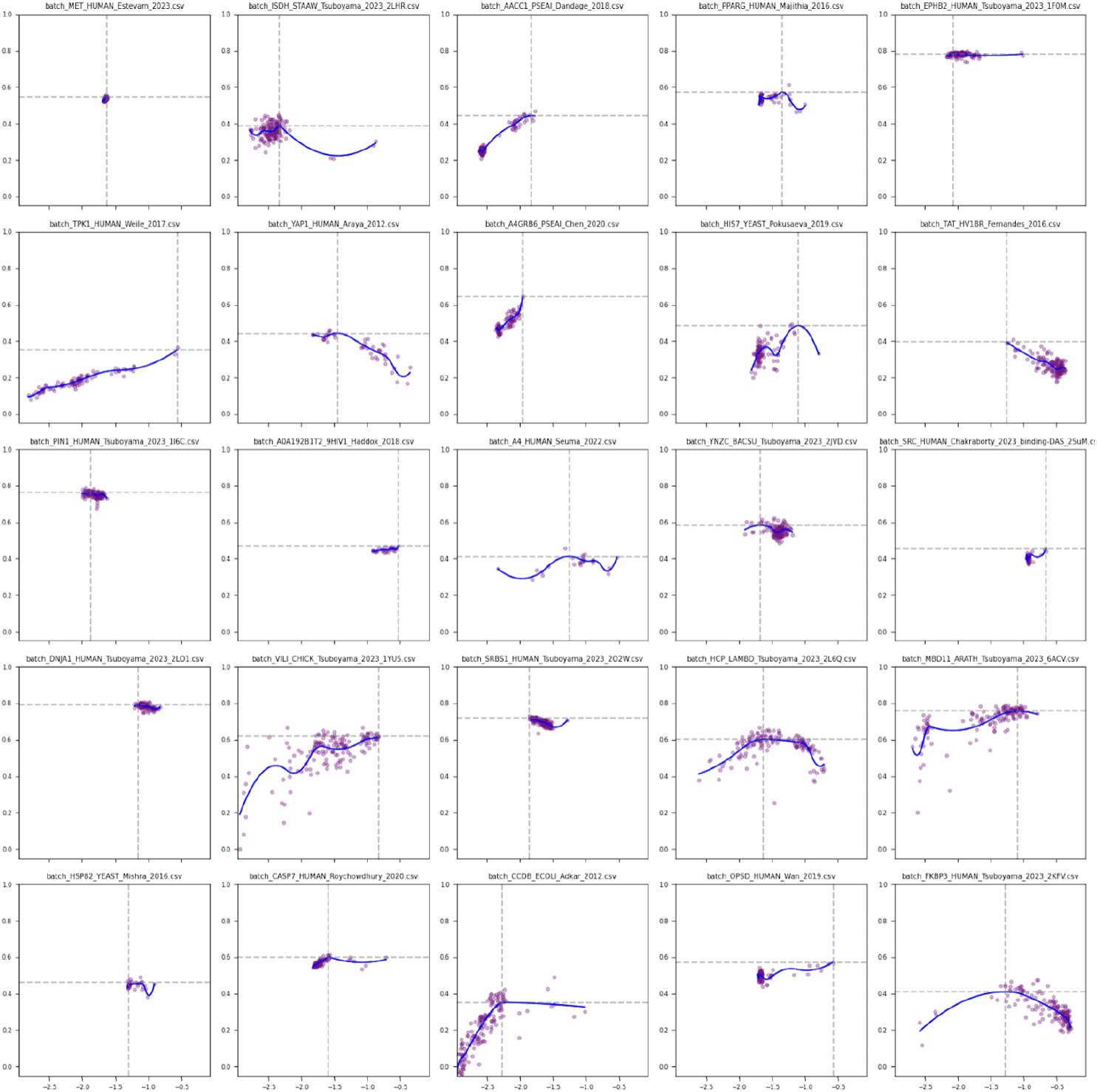
Log-likelihood and Spearman correlation for 25 randomly selected individual assays. Each point in the scatter plot is a different prompt (set of context sequences). Here we observe that several assays exhibit a decline in Spearman correlation once the log-likelihood exceeds a certain threshold.

### Limitations

While our in silico metrics are promising, we do not yet have experimental validation of the functionality of our generated sequences. Also, in our comparisons for sequence generation, we limited PoET’s inference to 8192 context tokens (matching ProFam-1’s training limit), although PoET is capable of processing longer contexts. Additionally, as PoET’s sequence sampling code is not open source, our re-implementation for these benchmarks may differ from the authors’ closed-source server/API. Finally, while competitive, ProFam-1 requires significantly more compute for zero-shot scoring (120 forward passes) to match the performance PoET achieves with fewer (15) passes.

By releasing ProFam-1, the ProFam Atlas, and our full training and inference codebase, we aim to provide a transparent and flexible foundation for the protein machine learning community. We hope this open resource will accelerate progress in developing models that not only capture evolutionary statistics but also robustly design functional proteins.

## 7 Availability

The complete ProFam codebase, including data preprocessing pipelines, training scripts, inference utilities for variant scoring and family-conditioned generation, is publicly available at https://github.com/alex-hh/profam/. ProFam-1 model weights are publicly accessible at https://huggingface.co/judewells/ProFam-1. The corresponding training data from the ProFam Atlas are available at https://zenodo.org/records/17713590.

## Acknowledgments

JW, WL, DM, NB, BP, CO, BR, MH acknowledge funding from the BBSRC grant ProtFunAI (BB/Y514044/1). JW acknowledges the receipt of studentship awards from the Health Data Research UK-The Alan Turing Institute Wellcome PhD Programme in Health Data Science (Grant Ref: 218529/Z/19/Z). JW was supported by the Encode AI for Science Fellowship. AHH was supported by the EPSRC Grant EP/S021566/1. This project was supported by the NVIDIA Academic Grant Program.

## A Data Sources

### A.1 FoldSeek AlphaFold Database Clusters

To include a structure-centric definition of protein family, we source data from the FoldSeek-based AlphaFold Database (AFDB) clusters [36]. These clusters were generated by applying first sequence-(50% sequence identity at 90% coverage) and then structure- (E-value<=0.01 at 90% coverage) based clustering to the AFDB, which resulted in 2.3 million non-singleton clusters. We treat each of these clusters as a protein family from which we can sample sequences to create a single training example, which we refer to as a *document*. For validation, we constructed a leakage-free set of 128 protein families and filtered the training data against it to ensure no sequence in any of the training datasets has above or equal 30 percent of pairwise identical residues (PID) against *any* sequence in one of the 128 held-out families (Appendix section B). After removing validation proteins, this set contains 2.3M protein families comprising 30M sequences.

### A.2 OpenProteinSet MSAs

The OpenProteinSet [26] is a widely used resource for defining protein families, notably for training structure prediction methods such as OpenFold [37]. This dataset originated from >16 million MSAs produced by HHblits aligning all-against-all on Uniclust30, from which the OpenFold authors filtered 270,000 maximally diverse MSAs. These MSAs may not be suitable for training protein family LMs, as sampling entries from the alignment can yield non-overlapping sequence segments with no pairwise similarity, or sequences with significant length differences. To remedy this and generate cleaner, length-consistent families, we re-processed the MSAs through a segmentation and clustering pipeline. We first split aligned sequences at contiguous gaps of *>* 10 residues, discarding resulting fragments shorter than 90 residues. The remaining subsequences were pooled and re-clustered using MMseqs2 [38] at 30% sequence identity and 70% coverage; clusters containing at least two sequences were defined as valid families. This pipeline yielded 37M protein families comprising 246M sequences.

### A.3 TED FunFams

While the FoldSeek AFDB clusters [39] provide a useful partitioning of the AFDB, sequences in this dataset are only retained if they achieve a conservative 90% overlap at the whole-chain level. To expand on this, we also considered homology at the domain level and incorporated alignments from fine-grained functional groupings. Specifically, we mapped all continuous domains from The Encyclopedia of Domains [25] that had a CATH assignment onto a library of 212,872 Hidden Markov Model profiles for the CATH Functional Families (v4.3.0) [40]. TED domain sequences were scanned with `hmmsearch` from the HMMER3 suite [41] using per-profile threshold cutoffs (`cut_tc`). Resulting matches were further refined with `cath-resolve-hits` [42] using default parameters to identify optimal domain boundaries. TED domains matching existing FunFam signatures were redundancy-reduced at 50% sequence identity, 90% overlap with MMseqs2 [43]. To ensure sequence and length consistency within these families, we clustered within families (30% sequence identity, 80% overlap with MMseqs2) and keep all members in the largest cluster only. This resulted in 38k families and 765k sequences.

### A.4 UniRef90 Single Sequences

To add coverage to regions of sequence space not covered by other family-based groupings, and to increase the model’s robustness in applications lacking a rich, explicitly provided, evolutionary context, we included 205M single sequences from UniRef90 (release 2025_01) [44]. Each document in the UniRef90 dataset consists of a single sequence.

#### Training data

| Dataset | Sample Probability | Max tokens per example | Families | Sequences |
| --- | --- | --- | --- | --- |
| FoldSeek S50 | 0.33 | 8192 | 2,300,407 | 30,033,967 |
| Uniref90 | 0.33 | 8192 | 203,929,771 | 203,929,771 |
| OpenFold | 0.33 | 320 | 37,245,744 | 246,251,236 |
| TED FunFams | 0.01 | 8192 | 38,097 | 765,136 |

## B Held-out Dataset Construction

We constructed the held-out validation dataset of 128 FoldSeek AFDB clusters and removed homologs from the training data based on sequence similarity:

1. **Select held-out families and remove them from training:** We selected 128 families from the FoldSeek clustering of the AlphaFold Database (AFDB) [24, 36, 39] and excluded all sequences belonging to these families from the candidate training pool.
2. **Build a non-redundant target set from held-out families:** We extracted all sequences from the 128 families and clustered them with MMseqs2 `easy-cluster` [38, 43] at 90% sequence identity and 80% alignment coverage. One representative per cluster was retained to form the held-out *target* database.
3. **Assemble the query set from all training sources:** We created a *query* database containing all remaining candidate training sequences across datasets (UniRef90, reprocessed OpenFold MSAs [26], and TED FunFams).
4. **Search and filter by homology:** We ran MMseqs2 `easy-search` from the query database against the held-out target database and removed any query sequence with a hit to any target representative at ≥30% sequence identity and ≥80% alignment coverage. The filtered remainder constitutes the final training set used in our experiments.

### B.1 EC Clustered Validation Dataset

EC family annotations were downloaded from https://ftp.expasy.org/databases/enzyme/enzyme.dat on 30 May 2024, and corresponding protein sequences were retrieved using the UniProt API. For each level-4 EC family, sequences were clustered with MMseqs2 at a minimum of 30% sequence identity and 70% coverage. To ensure family homogeneity, only sequences belonging to the largest cluster in each family were retained, and families whose largest cluster contained fewer than 50 sequences were discarded.

Sequences in each retained cluster were aligned with MAFFT (v7.526). The resulting MSAs were filtered using HHfilter (v3.3.0) with default parameters and a maximum sequence identity threshold of 90% to remove highly redundant sequences. Families with fewer than 50 sequences after filtering were removed, yielding a final set of 460 EC families.

These 460 families were used for sequence-based evaluation of first-order family statistics (Section C.2). For computation of statistical coupling analysis-style covariances (Section C.3), we restricted the analysis to the 24 families whose filtered MSAs contained at least 200 sequences, since deep MSAs are required to robustly estimate the second order statistics.

## C Benchmark Details

To evaluate ProFam-1’s capabilities in sequence generation and variant scoring, we utilized four distinct sets of benchmarks. First, we employed 128 held-out FoldSeek AFDB families to assess sequence generation quality and structural consistency. Second, we used Enzyme Commission (EC) families to evaluate the model’s ability to capture sequence statistics within functionally related groups; specifically, 460 EC families were used for first-order statistic evaluation and a subset of 24 deep families was used for Statistical Coupling Analysis (SCA). To test the utility of ProFam-generated synthetic MSAs, we benchmarked structure prediction performance on CASP 15 and 16 targets. Finally, we utilized the ProteinGym benchmark [31] for zero-shot variant scoring. Throughout these benchmarks, we extensively compare ProFam-1 against PoET [19], an autoregressive pfLM trained on sets of homologous sequences.

### C.1 Sequence Generation and Structural Consistency

To evaluate the structural plausibility of generated sequences, we utilized the 128 held-out families described in Section A.1. For each family, we sampled 20 sequences from ProFam-1 and PoET while conditioning on a random subset of the family’s sequences up to a maximum of 8192 tokens. For ProFam-1 in non-ensemble mode, we condition on the exact same prompt as PoET. For ProFam-1 in ensemble mode, we allow the prompts to differ. We selected the generated sequence with median length to predict its structure using ColabFold with default settings [23].

### C.2 Family Sequence Statistics

We ascertained whether ProFam-1 captures sequence statistics such as conservation and single-site variations from both minimal context (single sequences) and extended context (multiple sequences). For this evaluation, we utilized 460 Enzyme Commission (EC) families (construction details in Appendix B.1), chosen because they group proteins by shared catalytic function.

We used a single representative sequence per EC family to independently generate sequences using the sampling parameters described in Appendix E.1. We compared the per-position conservation correlation and KL divergence between the natural family MSA and the synthetic MSA generated by sampling sequences from PoET and ProFam-1. KL divergence is measured at each position in the MSAs, and we report the average. We further evaluated the model using multi-sequence prompts to assess how additional evolutionary context influences the fidelity of generated statistics. For these experiments, we define an acceptable length range for a generated sequence as being no less than 0.66 × shortest seq. in family and no greater than 1.5 × longest seq. in family.

### C.3 Higher-Order Statistics of Synthetic MSAs

First-order statistics derived from multiple sequence alignments (such as per-column conservation) are necessary but not sufficient to uniquely specify fold and activity [45]. Incorporating second-order statistics via pairwise couplings and associated statistical models (e.g. Statistical Coupling Analysis and Potts models [46, 47]) captures residue–residue constraints aligned with structural contacts [47], and enables the design of sequences that retain fold and function [48, 49]. To determine if ProFam-1 generated sequences correctly capture the pairwise couplings of protein families, we measure the level of correlation between residue pair covariances in synthetic MSAs and MSAs constructed from natural sequences, similar to [50, 51].

Let *L* be the aligned sequence length and *S* the alphabet size (here *S* = 20). For the natural and synthetic MSAs, we compute SCA-style rank 4 covariance tensors *C* where indices *i, j* index aligned positions ∈ {1, …, *L*} and indices *a, b* index the amino acid identities at positions *i* and *j* respectively:

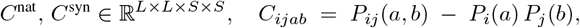

where probabilities are estimated from amino acid counts with a pseudocount of 1 added to each position, and gaps/unknowns excluded.

An entry (*i, j, a, b*) is kept if and only if all of the following hold:

1. Pair support filter: both MSAs have at least 10 sequences with non-gaps at positions *i* and *j*
2. *i* != *j*; (the diagonal is excluded)
3. Non-symmetric row filter at *i*: the count of residue *a* at position *i* is at least 10 in both MSAs; no residue-specific constraint is applied on *b* at position *j*.

We then calculate the Pearson correlation between the flattened covariance tensors, only for the retained entries. The covariance analysis was conducted on a subset of 24 deep EC families (from the 460 families described above) that possess sufficient homologs (>200 sequences) to accurately estimate covariance.

### C.4 Synthetic MSA Generation for CASP Targets

As co-variation strongly influences structure prediction, to demonstrate the utility of ProFam-1 we benchmark synthetic MSA generation to predict the structure of nine protein targets from CASP16 and 45 protein-domain targets from CASP15, comprising all targets that had resolved PDB structures at the time of writing. CASP16 predictions were made at the whole chain level, while CASP15 was predicted at the domain level, due to source data formatting. Towards this end, we provided ProFam-1 only with the target sequence (no additional family information) to generate 1200 sequences (sampled independently of each other) for each target protein in CASP15/16. After alignment of the generated sequences, the resulting MSA was input to ColabFold/AlphaFold2 [23]. For comparison, we also compute scores for structure predictions made without an MSA (single-sequence input) and for predictions using an MSA of natural homologs generated with the default ColabFold protocol.

### C.5 Zero-Shot Variant Fitness Prediction

Recent work has shown that the log-likelihood assigned to a variant sequence by a pLM is positively correlated with the variant’s empirical fitness across many functional assays [31]. This enables the use of pretrained pLMs as zero-shot fitness predictors, without requiring any assay-specific training data.

We benchmark protein-variant fitness prediction using the ProteinGym dataset [31], a collection of deep-mutational scanning (DMS) experiments spanning approximately 200 diverse proteins. For each protein, the dataset contains an experimentally derived fitness measurement for hundreds of variants across various functional assays, including thermal stability, enzymatic activity, and ligand binding. Computational methods are assessed based on the Spearman rank correlation between the predicted fitness score and the fitness assay result. To aid comparison with PoET [19], we used the same MSAs used by PoET.

## D Diversity Weighting for Prompt Sequence Sampling

To mitigate the risk that the prompt might sample highly similar sequences from a large cluster within the family, we employ a diversity weighting scheme taken from [19]. The homology-based weighting scheme assigns each sequence in a multiple sequence alignment (MSA) a weight inversely proportional to the number of similar sequences in the MSA.

Given an MSA with sequences 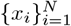, each of aligned length *L*, let the similarity between sequences *i* and *j* be

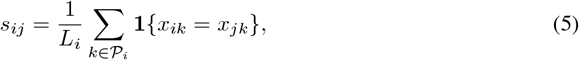

where 𝒫_*i*_ is the set of positions in sequence *i* that are not gaps or non-standard residues, and *L*_*i*_ = |𝒫_*i*_| is the number of such positions. The corresponding distance is

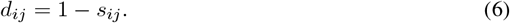

For a fixed threshold *θ* (in our case *θ* = 0.2), the neighbor count for sequence *i* is

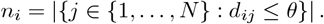

The unnormalised weight of sequence *i* is

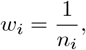

and the final normalised weight returned by the function is

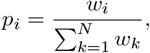

so that 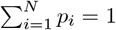.

## E Sequence Generation

### E.1 Sampling Hyperparameters

When sampling from sequences from PoET and ProFam-1, we limit the number of tokens in the prompt to the maximum sequence length seen by ProFam during training (8192 tokens) minus 1.2× the maximum sequence length in the prompt. This ensures that ProFam can generate a sequence that is 20% longer than any other sequence in the prompt while remaining in the supported token length range of 8192. We use a temperature of 1.0 for both ProFam-1 and PoET. ProFam-1 uses a top-p (nucleus sampling probability) of 0.95; PoET uses 0.9.

### E.2 Synthetic MSA Generation

In many of our benchmarks, we construct *synthetic* MSAs by iteratively sampling and subsequently aligning sequences from the model. We use these synthetic MSAs to assess whether the model captures the constraints of the protein family. This is done by comparing the synthetic MSA against the MSA of natural homologs using information-theoretic measures, such as per-position entropy-correlation, KL-Divergence, and SCA-style covariance. For both PoET and ProFam, sequences are generated independently using a fixed prompt or fixed ensemble of prompts (in the case of ProFam in ensemble mode). The diversity of generated sequences comes from the stochasticity of the autoregressive sampling process, rather than from conditioning on different evolutionary contexts. In order to compare corresponding columns in the natural MSA and the synthetic MSA, they must be aligned jointly. To achieve this, we concatenate the unaligned generated sequences to the unaligned natural sequences and align the resulting combined FASTA file with MAFFT (v7.526) using default settings.

## F Model Architecture and Hyperparameters

We used sequence packing [52] with Flash Attention [53], allowing multiple documents to be packed into a single batch dimension with no cross-document attention, thereby eliminating the need for any padding tokens. We used memory-mapped datasets to load the data efficiently and achieved a throughput of 212,000 tokens per second (53,000 tokens per GPU per second). The average number of tokens in a packed batch for a single GPU was 50,000 and we used data-parallelism with 4 GPUs and 2 gradient accumulation steps per update resulting in an effective batch size of 400,000 tokens per training step.

**Table 1:** Model architecture hyperparameter details.

| Field | Value |
| --- | --- |
| vocab_size | 68 |
| hidden_size | 1024 |
| intermediate_size | 4096 |
| num_hidden_layers | 16 |
| num_attention_heads | 16 |
| num_key_value_heads | 8 |
| max_position_embeddings (rope) | 131072 |
| rope_theta | 500000 |
| attn_implementation | flash_attention_2 |
| attention_bias | false |
| attention_dropout | 0.0 |
| rms_norm_eps | 1.0e-05 |
| hidden_act | silu |
| torch_dtype | bfloat16 |

**Table 2:**
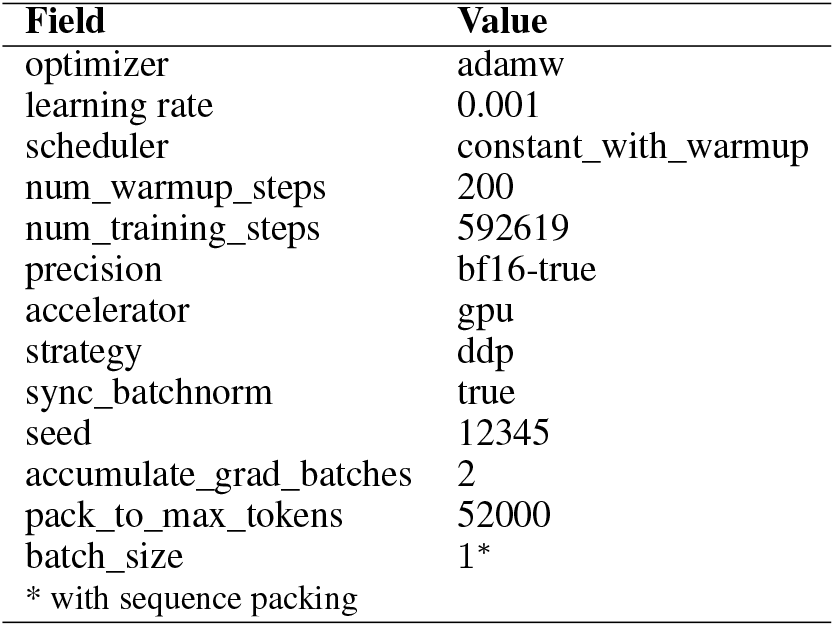
Training setup hyperparameter details.

| Field | Value |
| --- | --- |
| optimizer | adamw |
| learning rate | 0.001 |
| scheduler | constant_with_warmup |
| num_warmup_steps | 200 |
| num_training_steps | 592619 |
| precision | bf16-true |
| accelerator | gpu |
| strategy | ddp |
| sync_batchnorm | true |
| seed | 12345 |
| accumulate_grad_batches | 2 |
| pack_to_max_tokens | 52000 |
| batch_size | 1* |

\* with sequence packing

## G Additional Results

